# Lessening the bottleneck: reduced spatiotemporal overlap between krill fishing vessels and post-fledging chinstrap penguins led to increased apparent survival

**DOI:** 10.64898/2026.06.22.733719

**Authors:** Lucas Krüger, Francisco Santa Cruz, Magdalena Marquez, Juliana A. Vianna, Mercedes Santos, Andrea Piñones, César A. Cárdenas

## Abstract

Fledging is a critical period of a seabird life cycle. Using satellite telemetry, we compared movements and survival proxies (transmission duration) of chinstrap penguin fledglings tracked in 2017 (n=8) and 2025 (n=17) relative to krill fishing vessel activity. In 2017, fishing vessels operated intensively near colonies during summer, resulting in early, frequent encounters (median 1.3 days post-fledging) and short transmission durations (median 9.2 days). In 2025, reduced fishing delayed encounters (median 10.0 days) and tripled tracking duration (median 24.0 days). Hidden Markov Models revealed that vessel encounters reduced the probability of transitioning from foraging to transit behavior (β = -0.76), an effect stronger than the productivity (β = -0.11). While 87.5% of 2017 fledglings ceased transmission prematurely within weeks (half of those right after entering areas intensively used by fishing vessels), 65% of 2025 fledglings survived beyond March, with half of those five transmitting until May after dispersing eastward to the South Orkney Islands. These findings suggest that spatiotemporal overlap with krill fisheries during the critical post-fledging window affected foraging behavior and was associated with shorter transmission durations. Our results support further research of post-fledging penguin ecology to better understand the potential impact of fishery, and, following the precautionary principle, support fishing seasonal protection of important areas during critical periods of krill predators’ life cycle.

## Introduction

In the Antarctic Peninsula, chinstrap penguins (*Pygoscelis antarcticus*) are experiencing steep declines, likely attributable to multiple environmental change drivers affecting breeding success, recruit survival, and survival during the non-breeding season (Strycker et al., 2020; Talis et al., 2023). One of the most likely drivers is the availability of Antarctic krill (*Euphausia superba*), on which chinstrap penguins are heavily dependent during the breeding season, over 90% of the diet consists of Antarctic krill (Juáres et al., 2021; Panasiuk et al., 2020; Rozas Sia et al., 2026). Climate change is affecting Antarctic krill distribution, behavior, recruitment, and population dynamics through changes in sea ice extent, ocean temperature, and primary productivity (Atkinson et al., 2022; Kawaguchi et al., 2023), potentially increasing pressure on krill-dependent predators in affected regions.

A study found that fledgling mortality in the Antarctic Peninsula is highest during the first two weeks after fledglings leave the colony (Hinke et al., 2020). Fledging is a critical life-history stage that influence population dynamics and recruitment in birds (Ainley et al., 2018; Jones et al., 2026), and studies have suggested seabirds fledglings experience higher mortality when overlapping with fishing grounds (Afán et al., 2019; Ewbank et al., 2020). Juvenile survival is often lower than adults because fledglings must transition rapidly from parental dependence to independent foraging while facing environmental variability, predation risk, and energetic constraints in the marine environment (Jones et al., 2026; Swanson et al., 2023).

A potential additional source of interference is the Antarctic krill fishery, which has been operating in the Southern Ocean since the 1970s (Santa Cruz et al., 2022). The Commission for the Conservation of Antarctic Marine Living Resources (CCAMLR) was established in the 1980s to regulate fisheries in the Southern Ocean under an ecosystem-based management framework designed to minimize impacts on krill-dependent predators (Chavez-Molina et al., 2023; Constable, 2000; Miller and Slicer, 2014). CCAMLR also assumed the mandate of creating a network of Marine Protected Areas in its convention area (SC-CAMLR-XXIV) in order to safeguard representative marine ecosystems from unexpected synergistic effects of harvesting and climate change (see Brooks et al., 2020; Sylvester and Brooks, 2019; Teschke et al., 2021), for detailed discussions on the topic). In this regard, the Domain 1 Marine Protected Area (D1MPA) proposal, placed on the West Antarctic Peninsula and Southern Scotia Arc, the area where krill fishery has been concentrated in the last decades (Santa Cruz et al. 2022), among many conservation objectives, aims at providing precautionary measures by closing highly important areas for krill and dependent predators.

The objectives of this study were to compare chinstrap penguin post-fledgings movements in relation to the fishing fleet, while also testing for the effect of other likely influencing environmental drivers. We compared the birds tracked by Hinke et al. (2020) in 2017 with newly tracked fledglings from the 2025 fledging season. The temporal and spatial dynamics of the fishing fleet movements have changed in time to fish in the South Orkney Island surroundings (subarea 48.2) during late spring and summer, and moving to the Antarctic Peninsula and South Shetland Islands (subarea 48.1) in late summer and autumn (Santa Cruz et al. 2022). Therefore, the fishery operated extensively in subarea 48.1 during the 2017 breeding season, whereas fishing activity was substantially lower during the equivalent period in 2025. This contrast provides an opportunity to explore the potential effects of reduced fishery overlap on fledgling movements. We are testing the hypothesis that greater proximity to fishing vessels in 2017 was associated with earlier transmission failure, a proxy for mortality (i.e. Hinke et al. 2020) compared to 2025, in addition to the effects of environmental variables experienced by the birds. If supported, this hypothesis would suggest that reducing spatiotemporal overlap between fledging penguins and krill fishing vessels may benefit fledgling survival, and would provide information relevant to the evaluation of fishery management measures, including those proposed within the D1MPA framework.

## Methods

### Study area and penguin tracking

*Pygoscelis* penguins were tracked with Argos Platform Transmitter Terminals (PTTs) during the fledging seasons of 2016/2017 and 2024/2025 (hereafter 2017 and 2025 respectively). The 2017 data came from Hinke et al. (2020), who deployed transmitters on 8 chinstrap penguins, at two different sites (Cierva Cove 64.14°S, 60.98°W; Cape Shirreff 62.46°S, 60.79°W).

In the 2025 season, 20 chinstrap penguins in Kopaitic Island (Duroch Islands, 63.32°S, 57.92°W) were tagged with Spot tags configured to transmit 50 pings per hour. Tags were attached to dorsal feathers using marine-grade epoxy and cable ties. Each tagged penguin had a minimum body mass of 3 kg to minimize the potential additive effect of tagging on mortality. Tagging followed standard procedures and methods approved by ethical evaluation (Krüger et al., 2024b). Two penguins that transmitted just a few hours were excluded from the analysis, and one PTT did not send any data. Therefore, data from 25 penguins were used for the analysis (eight in 2017 and 17 in 2025).

### Tracking data processing

Only Argos positions classified as quality 1, 2, or 3 were retained, corresponding to estimated location errors of approximately 250 m, 500 m, and 1000 m, respectively. Tracking data was processed using a correlated random walk (CRW) approach implemented in the ‘*aniMotum*’ R package (Jonsen et al., 2023), fitted in 30-min intervals. Speed filters were used as, eventually, implausible locations are classified as high quality (Douglas et al., 2012). Model fit was validated by examining residuals of predicted locations (Supplementary Fig. S1). Predicted trips were rerouted to avoid unreal land crossings.

### Factors affecting fledgling movements and survival

Marine productivity is often associated with successful foraging in seabirds, as productivity serves as a proxy for food availability (Afán et al., 2014; Boersma et al., 2009; Ramírez et al., 2017). Primary productivity during summer is particularly important for Antarctic krill growth rates (Bahlburg et al., 2023; Mardones et al., 2026), recruitment rates, and abundance (Ryabov et al., 2023; Salmerón et al., 2023). Marine currents can act as a movement deterrent. Birds that must constantly swim against the current likely require more food to maintain high energy expenditure (Gunner et al., 2025).

Daily net primary production (NPP), expressed as grams of carbon per unit volume of seawater, was downloaded from the Copernicus Marine Services platform (https://data.marine.copernicus.eu/). We used the global ocean biogeochemistry analysis and forecast product (GLOBAL_MULTIYEAR_BGC_001_029). Daily Eastward and northward seawater velocity components were downloaded from the Copernicus Global Ocean Physics Reanalysis product (GLOBAL_MULTIYEAR_PHY_001_030).

Environmental variables were extracted at the penguins’ CRW-predicted tracking positions. Because tracking data were recorded hourly while environmental variables were available at a daily scale, the extracted values represent the average conditions each bird experienced on a daily basis.

### Krill fishery data

Fishing vessel tracks from the Automatic Identification System (AIS) were downloaded from the Global Fishing Watch platform for all vessels between January and May of each year (2017 and 2025), corresponding to the maximum duration of penguin tracking.

### Krill fishery proximity metrics

Subsequently, for each penguin location, we identified all fishing vessel positions within a ±30-minute time window and calculated the great-circle distance to each vessel. A proximity event was defined as any penguin position with at least one vessel within 50 km and within the 1⍰hour time window. A threshold of 50 km was selected to capture potential interactions occurring at spatial scales relevant to penguin movements and local fishery operations while accommodating location uncertainty associated with Argos telemetry.

### Behavioral state classification

To characterize the behavioral states of chinstrap penguin fledglings, we fitted a Hidden Markov Model (HMM) to the movement data. From the continuous-time correlated random walk predictions, we calculated step lengths and turning angles for each successive 30-minute interval. Step lengths (in kilometers) were computed using the Haversine formula using the ‘*geosphere*’ R package. Turning angles (in radians, range −π to π) were calculated as the direction change between consecutive steps and were subsequently wrapped to the interval [−π, π] using the atan2 function to ensure circular continuity.

Steps exceeding 20 km per 30 minutes (approximately 40 km/h, well above the maximum swimming speed of penguins) were considered unrealistic and removed from the dataset. This filtering affected less than 0.1% of all steps.

We fitted a two⍰state HMM to the step length and turning angle data using the ‘*momentuHMM*’ package in R. Step lengths were modeled using a gamma distribution, which is appropriate for continuous, positive-valued movement data. Turning angles were modeled using a von Mises distribution, a circular distribution suitable for angular data.

The HMM was parameterized with two behavioral states, which we a priori labeled as “Foraging” and “Transit” based on expected movement characteristics: foraging was expected to be associated with shorter step lengths and lower turning angle concentration, while transit was expected to be associated with longer step lengths and higher turning angle concentration (indicating directed, persistent movement).

Starting parameters for the HMM were informed by the empirical distributions of step lengths and turning angles. Initial step length means were set to 0.7 km (state 1) and 1.2 km (state 2), with standard deviations of 0.1 and 0.3, respectively. Initial turning angle concentrations were set to 0.01 (state 1) and 0.5 (state 2), reflecting low and moderate directional persistence. These values were chosen based on the 33^rd^ and 67^th^ percentiles of the observed step length distribution and visual inspection of the turning angle histogram.

We modeled the transition probabilities between behavioral states as functions of four covariates: (1) vessel encounter status (encounter; binary: TRUE if a penguin was within 50 km and 1 hour of an active fishing vessel), (2) net primary productivity, (3) current speed, and (4) minimum distance to the nearest vessel. All continuous covariates were scaled to have mean zero and unit standard deviation to facilitate model convergence and interpretation of coefficients. Missing values in current speed (0.08% locations) were replaced with zero (the mean after scaling). The model structure allowed us to test whether environmental conditions and vessel proximity influenced the likelihood of behavioral switching.

The HMM was fitted using maximum likelihood estimation via the fitHMM function. Following model fitting, the most probable state sequence for each individual was decoded using the Viterbi algorithm. State probabilities (smoothed estimates of the probability of being in each state at each time point) were extracted and subsequently used to calculate the mean probability of foraging per individual and the standard error of that estimate.

All analyses were conducted in R (R-Core-Team, 2026).

### Data availability statement

Tracking data for 2017 penguins is openly available in (Hinke et al., 2020), and tracking for 2025 penguins is openly available at (Krüger et al., 2026). Shapefiles of Antarctica were downloaded from the Antarctic Digital Database (Gerrish et al., 2026). Krill fishing vessels AIS were downloaded from the Global Fishing Watch map application (https://globalfishingwatch.org/map). R Codes are openly available at github repository (https://github.com/drlucaskruger/PenguinFledglings-KrillFishingVessels.git).

## Results

### Penguins movements

A total of 25 penguins were tracked across both breeding seasons, yielding 4,734 locations, which were interpolated to 27,065 locations after fitting CRWs. Median transmission duration was 18.9 days (75% IQR 7.8 to 27.9 days, maximum of 83 days). Median tracking duration in 2025 was almost threefold that of 2017: 24 days (IQR 10.7, 33.2) and 9.2 (IQR 7.7, 12.9) respectively.

In 2017, penguins from Cape Shirreff started moving west by 18 and 19 February before turning east within the Bransfield Strait, with the exception of one penguin that headed straight ahead to east in persistent movement after leaving the colony (fig 2a). That individual was the only penguin that continued transmitting beyond march and reached subarea 48.4 by April (fig 1a). Cierva Cove penguins started moving east by 26 to 28 February, but then remained within the Bransfield Strait until they stopped transmitting. None transmitted beyond march and none left subarea 48.1 (fig 1a).

**Figure 1.**
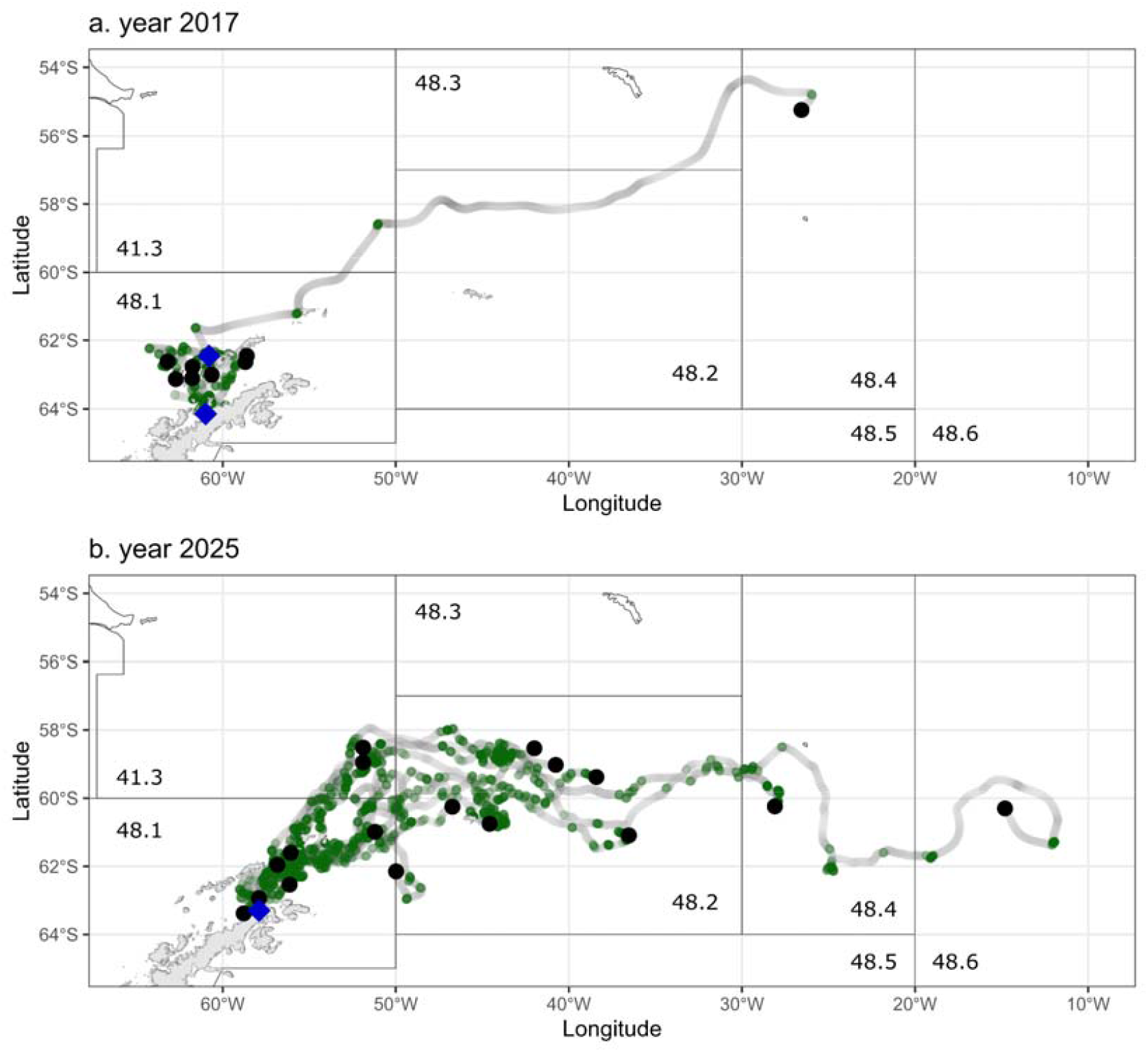
ARGOS tracking locations of chinstrap penguins (*Pygoscelis antarcticus*) fledglings in 2017 (a) and 2025 (b). Light gray points: all tracked positions. Dark green points: positions classified as foraging by HMM. Blue diamonds: colony locations (Cape Shirreff and Cierva Cove in 2017; Kopaitic Island in 2025). Black circles: last recorded position per individual. The name of the FAO fishing areas (spatial polygons) is presented within each subarea.

**Figure 2.**
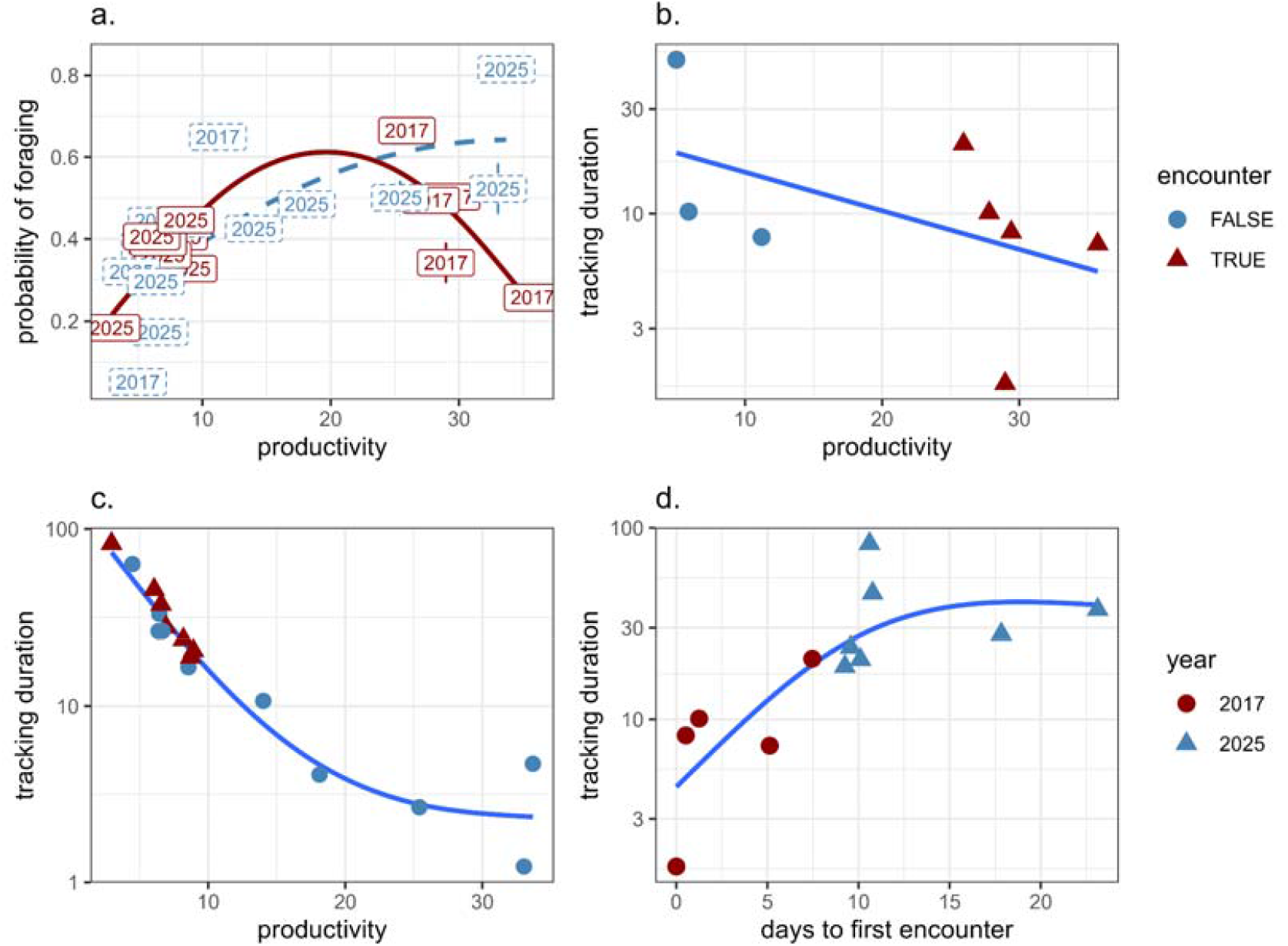
Individual-level relationships of chinstrap penguins (*Pygoscelis antarcticus*) fledged from three colonies in subarea 48.1 with productivity (NPP), vessel encounters, foraging, and survival. (a) Foraging probability vs. NPP: Vessel-encountering individuals (red triangles) non-encountering individuals (blue circles). (b) Tracking duration vs. NPP in 2017: Vessel-encountering individuals (red triangles) non-encountering individuals (blue circles). (c) Tracking duration vs. NPP in 2025: Vessel-encountering individuals (red triangles) non-encountering individuals (blue circles). (d) Timing of first vessel encounter vs. tracking duration: 2017 (red circles) 2025 (blue triangles). Points represent individual birds (n = 25); lines show fitted trends (GAM with k=4). Error bars in (a) indicate ±1 standard error.

Conversely, data from 2025 showed that penguins from Kopaitic Island had a more variable fledging date, from 28 February to 06 March. From the start they moved east, only 6 penguin stopped transmitting still within subarea 48.1, over half (11 out of 17) penguins kept transmitting through the second half of march, five penguins even transmitting until may, reaching subareas 48.2, 48.4 and even 48.6 (fig 1 b).

Marine productivity experienced by birds varied widely and was considerably higher in 2017 compared to 2025: 26.9 mg/m3 (9.9, 29.1) and 8.2 (6.4, 14.1) respectively. Median current speeds experienced by birds in 2025 were higher than in 2017: 0.38 (IQR 0.33, 0.44) in 2025 and 0.22 km/h (IQR 0.21, 0.33) in 2017.

Overall, 48% of penguins had encounter rates with vessels over zero (at least one vessel within 50 km and 1 hour of a fishing vessel). Encounter rates varied from 0% to 31.3%, over twice as much in 2017 (9.1% median, IQR 0%, 20.6%) in comparison to 2025 (0% median, IQR 0%, 1%). Days to first encounter after fledging were considerably shorter in 2017 (median 1.3 days; IQR 0.5 to 5.1) compared to 2025 (median 10 days; IQR 9.8 to 14.3). Penguins that encountered vessels had a median 12.3 km (IQR 6.0km, 19.7km) minimum distance in 2017 compared to 23.4 km (IQR 7.0km, 33.3 km) in 2025.

The Hidden Markov Model (HMM) fitted to chinstrap penguin movement data identified two distinct behavioral states, which we interpreted as foraging and transit based on step length and turning angle distributions. The foraging state was characterized by shorter step lengths (mean = 0.56 km per 30 min, SD = 0.42) and moderate turning angles (concentration = 13.1), consistent with area-restricted search behavior. In contrast, the transit state exhibited longer step lengths (mean = 1.24 km per 30 min, SD = 0.48) and highly directed movement (concentration = 666.9), indicative of commuting between areas.

At departure from the colony, fledglings showed a strong bias toward the transit state, with an initial probability of 0.91 (91%) for transit (table 1). This suggests that immediately after fledging, penguins prioritize dispersal. Behavioral states were highly persistent. The probability of remaining in the foraging state from one 30⍰min interval to the next was 0.93, while the probability of remaining in transit was 0.97. Consequently, switching from foraging to transit occurred with a probability of only 0.07 per 30 min, and switching from transit to foraging with a probability of 0.03 per 30 min. This high temporal autocorrelation indicates that individuals commit to a behavioral mode for extended periods, rather than switching frequently.

**Table 1.** Behavioral states transition coefficients for post-fledging chinstrap penguins (*Pygoscelis antarcticus*) in response to encounter with and distance to krill fishing vessels, marine primary productivity and marine currents, calculated using a Hidden Markov Model approach.

|  | Foraging | Transit |
| --- | --- | --- |
|  | Foraging -> Transit | Transit -> Foraging |
| Encounter | -0.76 | -0.01 |
| Distance | -0.05 | -0.3 |
| Productivity | -0.11 | 0.1 |
| Current Speed | 0.05 | -0.27 |

Vessel encounters had a strong negative effect on foraging-to-transit transition (β = -0.76), indicating that the presence of fishing vessels reduced the likelihood of transiting (table 1). In other words, penguins encountering vessels were less likely to transition into transit behavior, remaining longer in a given area. Distance to vessels showed a weak negative effect (β = -0.05), suggesting that closer proximity marginally decreased the probability of switching to transit.

Net primary productivity (NPP) also had a negative effect on foraging-to-transit transition (β = -0.11), meaning that in more productive areas, penguins were less likely to abandon foraging. Current speed showed a negligible positive effect (β = 0.05), with little influence on the foraging-to-transit transition.

For the transition from transit-to-foraging (i.e., the decision to stop traveling and start foraging), vessel encounters had a small effect (β = -0.01), indicating that encountering vessels did not influence the probability of transitioning into foraging. Distance to vessels showed a stronger negative effect (β = -0.30), suggesting that closer proximity to fishing vessels actually increased the likelihood of switching to foraging behavior (distance increases, transition to foraging decreases, so when a vessel is closer, the inverse happens).

Net primary productivity had a positive effect in transit-to-forage transition (β = 0.10), meaning that penguins were more likely to initiate foraging in areas with higher productivity, as expected. Current speed had a moderate negative effect (β = -0.27), indicating that stronger currents discouraged the transition into foraging.

At the individual level, the mean probability of foraging (derived from HMM state probabilities) showed a positive relationship with net primary productivity (Figure 2a). However, this relationship was non-linear and context-dependent. For individuals that encountered fishing vessels in 2017, the relationship followed a quadratic pattern, foraging probability increased with NPP at low to moderate productivity levels but declined at the highest productivity values. In contrast, individuals that never encountered vessels (including all 2025 individuals and non-encountering 2017 birds) showed a simpler positive relationship with no evidence of decline at high productivity. Importantly, these vessel-encountering individuals in 2017 also showed the shortest tracking durations (Figure 2b), contrasting with 2025 (figure 2c) which also had their first encounter with vessels later after fledging (figure 2d).

Those results are also consistent with the spatiotemporal distribution of encounters and last transmission (fig 3a, b). In 2017, all penguins from Cierva Cove colony stopped transmitting right after entering an area of high fishing effort (fig 3a). Movements indicated that penguins from Cierva Cove ceased transmission shortly after entering areas of high fishing activity, whereas most 2025 individuals continued moving eastward in the absence of intensive fishing activity near the colony (Fig 3b, Supplementary Video S1 and S2). In 2025, only two penguins stopped transmitting in areas of high fishing effort, by the South Orkney Island shelf in March (Fig 3b).

**Figure 3.**
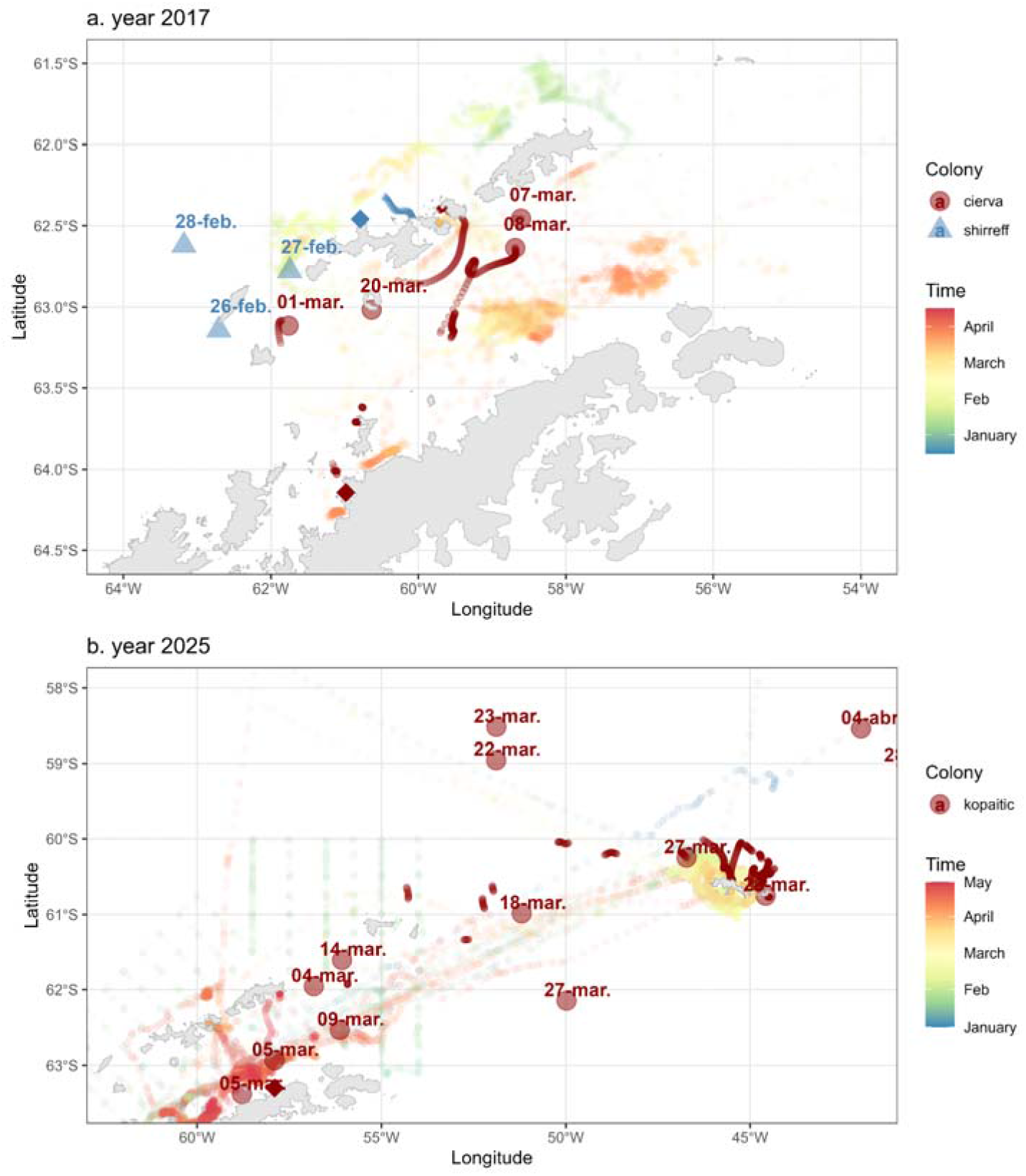
Temporal and spatial overlap between krill fishing vessels and chinstrap penguin (*Pygoscelis antarcticus*) fledglings in 2017 (a) and 2025 (b). Background light points: vessel positions colored by month (January to April). Foreground dark points: penguin last positions (small points) and encounter locations (large points), colored by colony of origin.

## Discussion

This study investigated whether spatiotemporal overlap between the Antarctic krill fishery and chinstrap penguin fledglings, comparing one year with fishing occurring consistently throughout summer (2017) and one year with less fishing during summer (2025). Our results reveal three main findings: (1) vessel encounters modified foraging probability, (2) the spatial pattern of encounters shifted from closer to the colonies within Bransfield Strait early overlap in 2017, to delayed eastward⍰shifted encounters in 2025 closer to South Orkneys Island, and (3) early first encounters substantially reduced duration of tag transmission. Together, these findings suggest that reducing spatiotemporal overlap between fledglings and fishing vessels during the critical early post-fledging period may improve apparent survival.

The combined results suggest that vessel encounters might interfere with the normal behavioral dynamics of chinstrap penguin fledglings. Specifically, encounters with fishing vessels reduced the probability of transitioning from foraging to transit, putting individuals in foraging mode for longer periods. This could reflect either competition forcing longer foraging bouts to meet energetic demands, or disturbance that disrupts normal movement decisions. Notably, vessel encounters *per se* did not influence the transition from transit to foraging, but the closer a vessel was, the more likely the penguins shifted from transit to foraging.

A well⍰documented example of fishery interfering with penguins comes from the African penguin (*Spheniscus demersus*), where fishery⍰induced ecological traps have been implicated in population declines (Sherley et al., 2017; Sydeman et al., 2021). In that system, adult penguins preferentially foraged in areas with high fish abundance, but those same areas attracted competing fisheries, leading to reduced survival and recruitment. Our results suggest a similar mechanism may operate for *Pygoscelis* penguins in the western Antarctic Peninsula: areas of high primary productivity (and thus high krill density) historically supported successful foraging and fledging, but these same areas attracted intensive krill fishing during summer. In 2017, penguins that foraged in productive nearshore waters experienced high vessel encounter rates and lower apparent survival. By 2025, fishing vessels had used those areas only in a shorter period of time due sea-ice conditions in subarea 48.2 (Supplementary video S3), and upstream the Kopaitic Island colony. Therefore, penguins that dispersed eastward encountered vessels only later.

Those levels of early transmission failure (seven out of eight in 2017) are also consistent with studies of fledglings encountering intensively fished areas after colony departure for other seabird species in the Mediterranean (Afán et al., 2019). Incidental mortality in Antarctic Krill fishery is considerably low due to CCAMLR strict conservation measures, and chinstrap penguins have been considered of low risk to incidental mortality (Crawford et al., 2017), appearing as incidental mortality in krill fishery at low numbers when CCAMLR reopened the working group in Incidental Mortality (WG-IMAF) in 2022 (CCAMLR Secretariat, 2023, 2022). Prior to that WG-IMAF was dedicated solely to longline fishery. When reviewing meeting reports, we were not able to find any mention of penguin incidental mortalities in krill fishery for previous years, nor find documented cases of chinstrap penguin incidental mortality in krill fishing nets in the CCAMLR Fishery Reports.

Fisheries might affect penguins through displacement, disrupting krill swarms (causing less dense or deeper swarms), or by disrupting the behavior of the penguin itself. Although effects of noise pollution are well documented for cetaceans, that is an area less studied for penguins (but see Ainley and Wilson, 2023; Pichegru et al., 2022). Interactions with fishing nets and gear, in analogy to the quantification of warp-strikes of flying seabirds, also are inexistent and unlikely to be quantified.

We interpreted that premature tag transmission cessation as a proxy, but tags could also fail due to battery depletion, antenna damage, or detachment (Kooyman et al., 2015). However, the rapid and early cessation in 2017 (many within 10 days) is more consistent with mortality than with tag failure, given that tags were rated for 60 to 90 days of operation (Hinke et al. 2020). Penguin fledglings in other areas of Antarctica transmitted on average for much more days than the ones in 2017 (Clarke et al., 2003; Makhado et al., 2025).

The 2017 birds originated from different colonies than the 2025 birds. Although all are located in the same region, differences in local oceanography or krill availability could have influenced apparent survival independently of year effects. Gerlache Strait was one the most heavily fished areas in 2025, with some of that catch being caught during early summer (Supplementary Video S3), before fledging. But there was no tracking of fledglings in 2025 in Cierva Cove colony to compare. Future studies should track birds from multiple colonies within the same year to disentangle colonies⍰specific effects from temporal changes in fishery behavior. In addition, we used NPP as a proxy for krill habitat, but primary productivity does not always translate directly to krill biomass or accessibility to penguins. Direct measurements of krill abundance (e.g., from acoustic surveys) would strengthen future studies.

### Are predators also targeting these same areas?

It is worth noting that natural predators, including leopard seals (*Hydrurga leptonyx*) and killer whales, also concentrate in productive nearshore waters during summer (Fearnbach et al., 2019; Kienle et al., 2021; Krause et al., 2016; Pitman and Durban, 2010). Thus, even in the absence of fishing, fledglings face elevated predation risk in these habitats. However, the additive effect of fishery interference (e.g., movement disruption, competition for krill) may push mortality up.

### Fishery overlap and apparent survival

Spatial encounter maps provide a mechanistic explanation for the differences found between years. In 2017, fishing vessels operated intensively in the Bransfield and Gerlache straits, overlapping directly with fledgling departures from colonies at Shirreff and Cierva Cove. Encounters occurred immediately and continuously, coinciding with shorter transmission durations and reduced eastward dispersal (Supplementary Video S1). In 2025, vessels largely avoided the Peninsula until late March. Consequently, penguins departing from Kopaitic Island colony encountered no vessels in nearshore waters; only those that dispersed eastward toward the South Orkney Islands eventually overlapped with the fleet, and these encounters occurred later in the tracking period (Supplementary Video S2).

Chinstrap penguins have been declining consistently in the subarea 48.1 and moderately in 48.2 (Talis et al., 2023). Preliminary evidence suggests that even in the larger colonies considered stable or increasing until 10 years ago (Lynch et al., 2016) the trend now is reversing (Gregory and Belchier, 2026; Ratcliffe et al., 2025), raising conservation concerns towards protection of this species.

### Implications for MPA design and fishery management

Our findings have direct relevance for the proposed Domain 1 Marine Protected Area D1MPA and for the precautionary management of the Antarctic krill fishery under CCAMLR. Survival during the first weeks post-fledging is a sensitive and tractable metric for assessing fishery interference. Monitoring programs within the D1MPA could compare survival of fledglings from colonies inside vs. outside the proposed MPA boundaries, ideally before and after full implementation of seasonal closures. Such comparisons would provide direct evidence of MPA benefits and support adaptive management. Our results support further evaluation of seasonal fishing closures (e.g., November to March) during periods when fledging penguins are most vulnerable to overlap with fishing activity. Noteworthy that other life stages, like post-breeding adults, could also benefit from such a temporal closure in specific periods of time. Therefore formalizing protection zones (year-round or seasonal, Krüger et al., 2024a) through mechanisms such as those proposed through the D1MPA, could provide a framework for long⍰term monitoring and protection at critical stages for predators.

## Conclusions

This study provides evidence that spatiotemporal overlap between the Antarctic krill fishery and fledging penguins was associated with shorter transmission durations (a proxy for apparent survival) in 2017. A reduction of fishing activity during 2025, could have led to reduced overlap and potentially increased fledging apparent survival. Our results support that reducing seasonal spatiotemporal overlap may reduce interference with fledging penguins during the critical first weeks after departure.

## Supplementary files legends

Supplementary figure S1. Diagnostic plots for the continuous□time correlated random walk (CTCRW) model fitted to chinstrap penguin tracking data. Panels a-d show residuals over time for eastward (x) and northward (y) coordinates, separately for 2017 and 2025. Panels e-f are quantile□quantile (QQ) plots comparing residuals to a normal distribution. Panels g-h show autocorrelation functions (ACF) for x and y residuals. Dashed lines represent zero (a-d), theoretical normality (e-f), or 95% confidence intervals (g-h). Smooth lines (a-d) are local regressions (LOESS). Grey points indicate individual residuals.

Supplementary Videos S1 and S2. Animated maps showing daily positions of *Pygoscelis* penguin fledglings (red circles) and krill fishing vessels (grey triangles) during the austral summers of 2017 (S1) and 2025 (S2). Blue diamonds indicate breeding colony locations: Shirreff Cove and Cierva Cove (2017); Kopaitic Island (2025). Grey landmass represents the Antarctic coastline (Antarctic Digital Database). Penguin positions were interpolated from Argos tracks using a continuous□time correlated random walk model (CRW; aniMotum package). Vessel positions were obtained from Global Fishing Watch AIS data. The animations illustrate the temporal and spatial overlap between dispersing fledglings and active fishing grounds, with 2025 showing a decreased encounter frequency compared to 2017.

Supplementary Video S3 Time series of sea ice concentration (SIC) from December 1, 2024, to January 31, 2025. Colors represent daily sea ice concentration ranging from 0 (white, ice-free) to 1 (dark blue, fully ice-covered). Blue triangles mark penguin colonies at Shirreff, Cierva, and Kopaitic. Vessel tracks are shown by colored symbols, with different colors representing individual ship IDs. The animation demonstrates the temporal dynamics of ice coverage during the late spring and early summer that prevented fishery from using the South Orkney Island fishing hotspot. Only vessels that fished in December 2024 and January 2025 are presented.

## References

Afán, I., Navarro, J., Cardador, L., Ramírez, F., Kato, A., Rodríguez, B., Ropert-Coudert, Y., Forero, M.G., 2014. Foraging movements and habitat niche of two closely related seabirds breeding in sympatry. Mar. Biol. 161, 657–668. 10.1007/s00227-013-2368-4

Afán, I., Navarro, J., Grémillet, D., Coll, M., Forero, M.G., 2019. Maiden voyage into death: are fisheries affecting seabird juvenile survival during the first days at sea? R. Soc. Open Sci. 6, 181151. 10.1098/rsos.181151

Ainley, D., Dugger, K., La Mesa, M., Ballard, G., Barton, K., Jennings, S., Karl, B., Lescroël, A., Lyver, P., Schmidt, A., Wilson, P., 2018. Post-fledging survival of Adélie penguins at multiple colonies: chicks raised on fish do well. Mar. Ecol. Prog. Ser. 601, 239–251. 10.3354/meps12687

Ainley, D.G., Wilson, R.P., 2023. Penguins Coping with a Changing Ocean, in: The Aquatic World of Penguins, Fascinating Life Sciences. Springer International Publishing, Cham, pp. 437–458. 10.1007/978-3-031-33990-5_13

Atkinson, A., Hill, S.L., Reiss, C.S., Pakhomov, E.A., Beaugrand, G., Tarling, G.A., Yang, G., Steinberg, D.K., Schmidt, K., Edwards, M., Rombolá, E., Perry, F.A., 2022. Stepping stones towards Antarctica: Switch to southern spawning grounds explains an abrupt range shift in krill. Glob. Change Biol. 28, 1359–1375. 10.1111/gcb.16009

Bahlburg, D., Thorpe, S.E., Meyer, B., Berger, U., Murphy, E.J., 2023. An intercomparison of models predicting growth of Antarctic krill (Euphausia superba): The importance of recognizing model specificity. PLOS ONE 18, e0286036. 10.1371/journal.pone.0286036

Boersma, P.D., Rebstock, G.A., Frere, E., Moore, S.E., 2009. Following the fish: penguins and productivity in the South Atlantic. Ecol. Monogr. 79, 59–76. 10.1890/06-0419.1

Brooks, C.M., Chown, S.L., Douglass, L.L., Raymond, B.P., Shaw, J.D., Sylvester, Z.T., Torrens, C.L., 2020. Progress towards a representative network of Southern Ocean protected areas. PLOS ONE 15, e0231361. 10.1371/journal.pone.0231361

CCAMLR Secretariat, 2023. Summary of Incidental Mortality Associated with Fishing activities data collected during the 2023 season, and a draft method for the extrapolation of IMAF and warp strikes.

CCAMLR Secretariat, 2022. Summary of incidental mortality associated with fishing activities during the 2022 season, and review of incidental mortality data and warp strike data since 2012.

Chavez-Molina, Vasco., Nocito, E.S., Carr, E., Cavanagh, R.D., Sylvester, Z., Becker, S.L., Dorman, D.D., Wallace, B., White, C., Brooks, C.M., 2023. Managing for climate resilient fisheries: Applications to the Southern Ocean. Ocean Coast. Manag. 239, 106580. 10.1016/j.ocecoaman.2023.106580

Clarke, J., Kerry, K., Fowler, C., Lawless, R., Eberhard, S., Murphy, R., 2003. Post-fledging and winter migration of Adélie penguins Pygoscelis adeliae in the Mawson region of East Antarctica. Mar. Ecol. Prog. Ser. 248, 267–278. 10.3354/meps248267

Constable, A., 2000. Managing fisheries to conserve the Antarctic marine ecosystem: practical implementation of the Convention on the Conservation of Antarctic Marine Living Resources (CCAMLR). ICES J. Mar. Sci. 57, 778–791. 10.1006/jmsc.2000.0725

Crawford, R., Ellenberg, U., Frere, E., Hagen, C., Baird, K., Brewin, P., Crofts, S., Glass, J., Mattern, T., Pompert, J., Ross, K., Kemper, J., Ludynia, K., Sherley, R.B., Steinfurth, A., Suazo, C.G., Yorio, P., Tamini, L., Mangel, J.C., Bugoni, L., Uzcátegui, G.J., Simeone, A., Luna-Jorquera, G., Gandini, P., Woehler, E.J., Pütz, K., Dann, P., Chiaradia, A., Small, C., 2017. Tangled and drowned: A global review of penguin bycatch in fisheries. Endanger. Species Res. 34, 373–396. 10.3354/esr00869

Douglas, D.C., Weinzierl, R., C. Davidson, S., Kays, R., Wikelski, M., Bohrer, G., 2012. Moderating A rgos location errors in animal tracking data. Methods Ecol. Evol. 3, 999–1007. 10.1111/j.2041-210X.2012.00245.x

Ewbank, A.C., Sacristán, C., Costa-Silva, S., Antonelli, M., Lorenço, J.R., Nogueira, G.A., Ebert, M.B., Kolesnikovas, C.K.M., Catão-Dias, J.L., 2020. Postmortem findings in Magellanic penguins (Spheniscus magellanicus) caught in a drift gillnet. BMC Vet. Res. 16, 153. 10.1186/s12917-020-02363-x

Fearnbach, H., Durban, J.W., Ellifrit, D.K., Pitman, R.L., 2019. Abundance of Type A killer whales (Orcinus orca) in the coastal waters off the western Antarctic Peninsula. Polar Biol. 42, 1477–1488. 10.1007/s00300-019-02534-z

Gerrish, L., Ireland, L., Fretwell, P., Cooper, P., Skachkova, A., 2026. Medium resolution vector polylines of the Antarctic coastline - VERSION 7.12. 10.5285/37AD06FF-888D-452C-BE85-018E59600103

Gregory, S., Belchier, M., 2026. South Georgia & South Sandwich Islands Marine Protected Area Management Plan. 10.5281/ZENODO.18629695

Gunner, R.M., Quintana, F., Tonini, M.H., Holton, M.D., Yoda, K., Crofoot, M.C., Wilson, R.P., 2025. Penguins exploit tidal currents for efficient navigation and opportunistic foraging. PLOS Biol. 23, e3002981. 10.1371/journal.pbio.3002981

Hinke, J.T., Watters, G.M., Reiss, C.S., Santora, J.A., Santos, M.M., 2020. Acute bottlenecks to the survival of juvenile Pygoscelis penguins occur immediately after fledging: Acute bottleneck to fledgling survival. Biol. Lett. 16, 0–5. 10.1098/rsbl.2020.0645

Jones, T.M., Kaiser, S.A., Sillett, T.S., 2026. Post-fledging ecology of birds: emergent patterns, knowledge gaps, and future frontiers. Biol. Rev. 101, 237–267. 10.1111/brv.70080

Jonsen, I.D., Grecian, W.J., Phillips, L., Carroll, G., McMahon, C., Harcourt, R.G., Hindell, M.A., Patterson, T.A., 2023. aniMotum, an R package for animal movement data: Rapid quality control, behavioural estimation and simulation. Methods Ecol. Evol. 14, 806–816. 10.1111/2041-210X.14060

Juáres, M.A., Grech, M.G., Casaux, R., Negrete, J., Fógel, J., Coria, N.R., Santos, M.M., 2021. Size structure of Antarctic krill inferred from samples of Pygoscelid penguin diets and those collected by the commercial krill fishery. Mar. Biol. 168, 22. 10.1007/s00227-021-03831-0

Kawaguchi, S., Atkinson, A., Bahlburg, D., Bernard, K.S., Cavan, E.L., Cox, M.J., Hill, S.L., Meyer, B., Veytia, D., 2023. Climate change impacts on Antarctic krill behaviour and population dynamics. Nat. Rev. Earth Environ. 10.1038/s43017-023-00504-y

Kienle, S., Trumble, S., Kanatous, S., Goebel, M., Borras-Chavez, R., Costa, D., 2021. Phenotypic plasticity in the movement patterns, dive behavior, and diet of an Antarctic apex predator, the leopard seal. Biol. Mar. Mamm. 10.3389/fmars.2022.976019

Kooyman, G.L., McDonald, B.I., Goetz, K.T., 2015. Why do satellite transmitters on emperor penguins stop transmitting? Anim. Biotelemetry 3. 10.1186/s40317-015-0091-2

Krause, D.J., Goebel, M.E., Marshall, G.J., Abernathy, K., 2016. Summer diving and haul-out behavior of leopard seals (Hydrurga leptonyx) near mesopredator breeding colonies at Livingston Island, Antarctic Peninsula. Mar. Mammal Sci. 32, 839–867. 10.1111/mms.12309

Krüger, L., Deregibus, D., Rebolledo, L., Santa Cruz, F., Santos, M., 2024a. Modifying General Protection Zones and introducing Seasonal Protection Zones for harmonising the D1MPA and the fishery in Domain 1.

Krüger, L., Marquez, M., Santa Cruz, F., Vianna, J., 2026. Platform Terminal Transmitter data for Chinstrap Penguin (Pygoscelis antarcticus) fledglings from a colony in the Antarctic Peninsula. 10.5281/ZENODO.20309350

Krüger, L., Vianna, J.A., Cárdenas, C.A., 2024b. General methodology for fieldwork with seabirds to monitor Antarctic and Subantarctic ecosystems and Ethical Certifications. Zenodo 10.5281/zenodo.10640095

Lynch, H.J., White, R., Naveen, R., Black, A., Meixler, M.S., Fagan, W.F., 2016. In stark contrast to widespread declines along the Scotia Arc, a survey of the South Sandwich Islands finds a robust seabird community. Polar Biol. 39, 1615–1625. 10.1007/s00300-015-1886-6

Makhado, A.B., Dyer, B.M., Masotla, M., Lowther, A., 2025. Dispersal and habitat preference of juvenile emperor penguins —implications for conservation management.

Mardones, M., Jarvis Mason, E., Santa Cruz, F., Watters, G., Cárdenas, C., 2026. Disparate estimates of intrinsic productivity for Antarctic krill across small spatial scales under a rapidly changing ocean. PLOS One 21, e0322671. 10.1371/journal.pone.0322671

Miller, D., Slicer, N.M., 2014. CCAMLR and Antarctic conservation: The leader to follow?, in: Garcia, S.M., Rice, J., Charles, A. (Eds.), Governance of Marine Fisheries and Biodiversity Conservation. Wiley, pp. 253–270. 10.1002/9781118392607.ch18

Panasiuk, A., Wawrzynek-Borejko, J., Musiał, A., Korczak-Abshire, M., 2020. Pygoscelis penguin diets on King George Island, South Shetland Islands, with a special focus on the krill Euphausia superba. Antarct. Sci. 32, 21–28. 10.1017/S0954102019000543

Pichegru, L., Vibert, L., Thiebault, A., Charrier, I., Stander, N., Ludynia, K., Lewis, M., Carpenter-Kling, T., McInnes, A., 2022. Maritime traffic trends around the southern tip of Africa – Did marine noise pollution contribute to the local penguins’ collapse? Sci. Total Environ. 849, 157878. 10.1016/j.scitotenv.2022.157878

Pitman, R.L., Durban, J.W., 2010. Killer whale predation on penguins in Antarctica. Polar Biol. 33, 1589–1594. 10.1007/s00300-010-0853-5

R Core Team. 2026. R: A language and environment for statistical computing (Version 4.4.2]) [Computer software]. R Foundation for Statistical Computing, Vienna, Austria. https://www.R-project.org/

Ramírez, F., Tarroux, A., Hovinen, J., Navarro, J., Afán, I., Forero, M.G., Descamps, S., 2017. Sea ice phenology and primary productivity pulses shape breeding success in Arctic seabirds. Sci. Rep. 7, 4500. 10.1038/s41598-017-04775-6

Ratcliffe, N., Colesie, C., Convey, P., Hart, T., Cugniere, L., Clucas, G., Fretwell, P., Richter, N., Dickens, J., Fenney, N., Walters, M., Putz, K., Belchier, M., Gregory, S., Black, J., 2025. Darwin Plus Main & Streategic FIanl report: Evidence-based conservation of key biodiversity in the South Sandwich Islands.

Rozas Sia, M.G., Farace Rey, A., Soutullo, Á., Machado-Gaye, A.L., Negrete, J., Santos, M.M., Juáres, M.A., 2026. No need to travel far, krill takeaway: Pygoscelid penguins as ecological indicators through foraging trip distribution and diet in Danco Coast. Mar. Biol. 173, 101. 10.1007/s00227-026-04871-0

Ryabov, A., Berger, U., Blasius, B., Meyer, B., 2023. Driving forces of Antarctic krill abundance. Sci. Adv. 9, eadh4584. 10.1126/sciadv.adh4584

Salmerón, N., Belle, S., Cruz, F.S., Alegria, N., Finger, J.V.G., Corá, D.H., Petry, M.V., Hernández, C., Cárdenas, C.A., Krüger, L., 2023. Contrasting environmental conditions precluded lower availability of Antarctic krill affecting breeding chinstrap penguins in the Antarctic Peninsula. Sci. Rep. 13, 5265. 10.1038/s41598-023-32352-7

Santa Cruz, F., Krüger, L., Cárdenas, C.A., 2022. Spatial and temporal catch concentrations for Antarctic krill: Implications for fishing performance and precautionary management in the Southern Ocean. Ocean Coast. Manag. 223, 106146. 10.1016/j.ocecoaman.2022.106146

Sherley, R.B., Ludynia, K., Dyer, B.M., Lamont, T., Makhado, A.B., Roux, J.P., Scales, K.L., Underhill, L.G., Votier, S.C., 2017. Metapopulation Tracking Juvenile Penguins Reveals an Ecosystem-wide Ecological Trap. Curr. Biol. 27, 563–568. 10.1016/j.cub.2016.12.054

Strycker, N., Wethington, M., Borowicz, A., Forrest, S., Witharana, C., Hart, T., Lynch, H.J., 2020. A global population assessment of the Chinstrap penguin (Pygoscelis antarctica). Sci. Rep. 10, 1–11. 10.1038/s41598-020-76479-3

Swanson, N., Vaughan, N., Belling, N., Roman, L., 2023. Post-fledging survival of wedge-tailed shearwaters is linked to pre-fledge mass, which has decreased over 40 years. Mar. Ecol. 44, e12776. 10.1111/maec.12776

Sydeman, W.J., Hunt, G.L., Pikitch, E.K., Parrish, J.K., Piatt, J.F., Boersma, P.D., Kaufman, L., Anderson, D.W., Thompson, S.A., Sherley, R.B., 2021. South Africa’s experimental fisheries closures and recovery of the endangered African penguin. ICES J. Mar. Sci. 78, 3538–3543. 10.1093/icesjms/fsab231

Sylvester, Z.T., Brooks, C.M., 2019. Protecting Antarctica through Co-production of actionable science: Lessons from the CCAMLR marine protected area process. Mar. Policy 103720. 10.1016/j.marpol.2019.103720

Talis, E.J., Che-Castaldo, C., Hart, T., McRae, L., Lynch, H.J., 2023. Penguindex: a Living Planet Index for Pygoscelis species penguins identifies key eras of population change. Polar Biol. 10.1007/s00300-023-03148-2

Teschke, K., Brtnik, P., Hain, S., Herata, H., Liebschner, A., Pehlke, H., Brey, T., 2021. Planning marine protected areas under the CCAMLR regime – The case of the Weddell Sea (Antarctica). Mar. Policy 124, 104370. 10.1016/j.marpol.2020.104370

