## Supplementary figures and images for "Lessening the bottleneck: reduced spatiotemporal overlap between krill fishing vessels and post-fledging chinstrap penguins led to increased apparent survival"

### supplementary figure S1

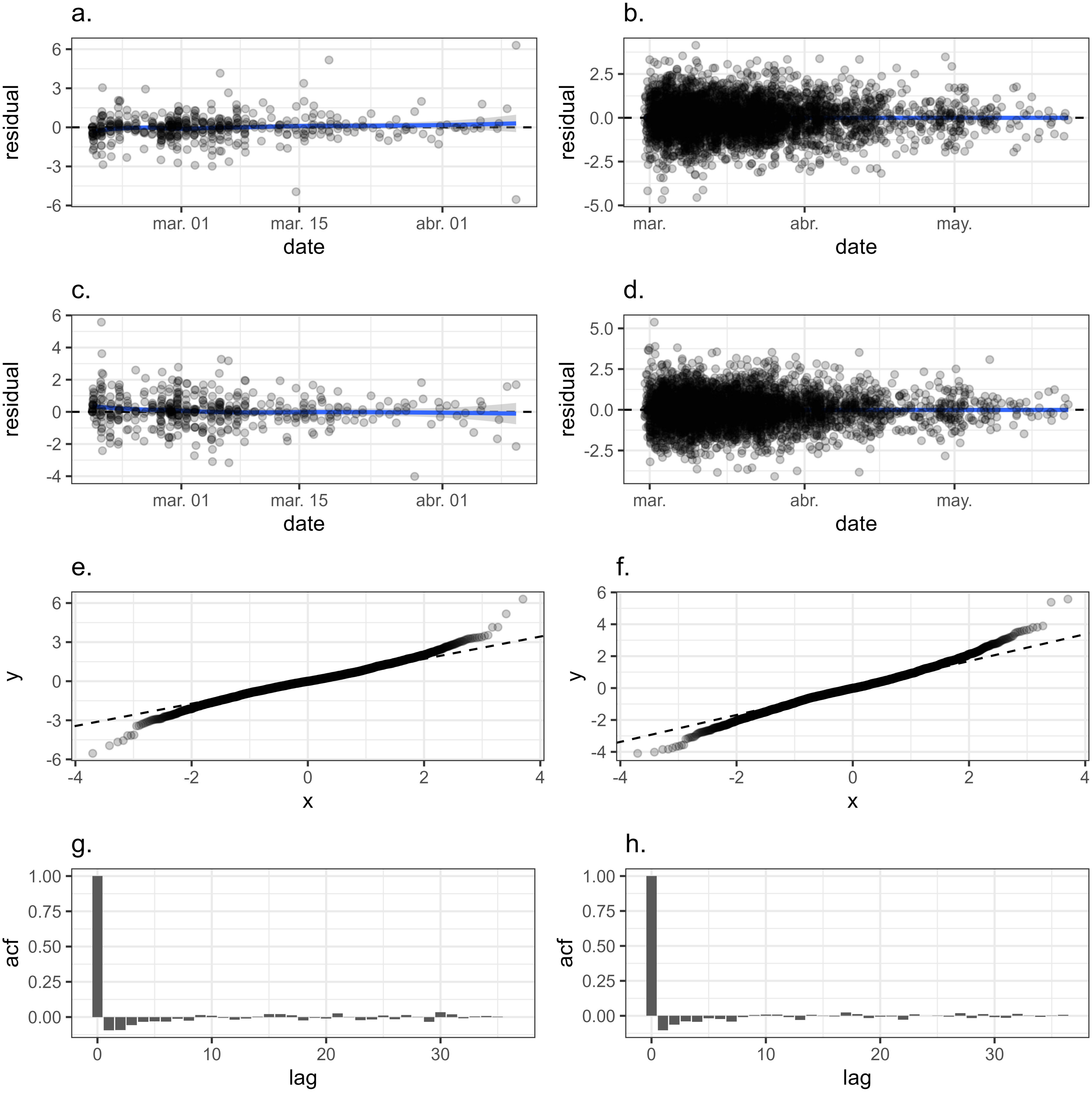

### supplementary video S1

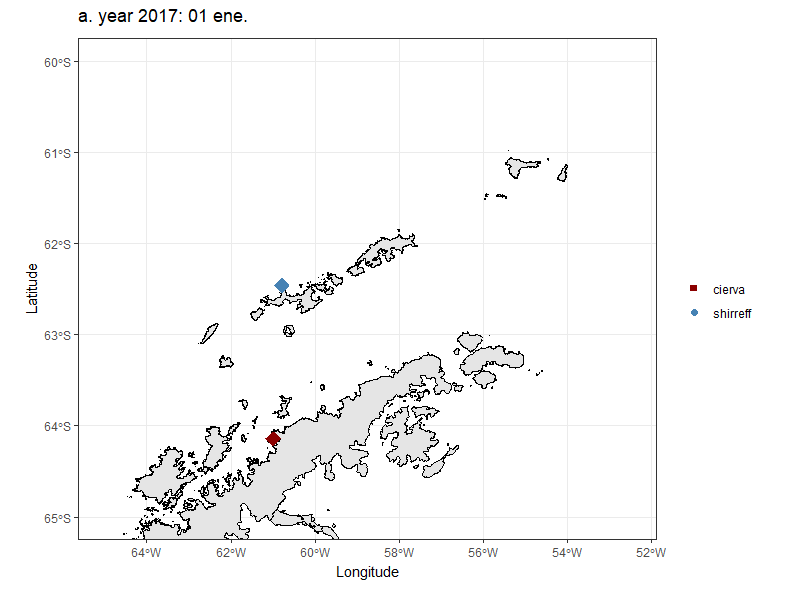

### supplementary video S2

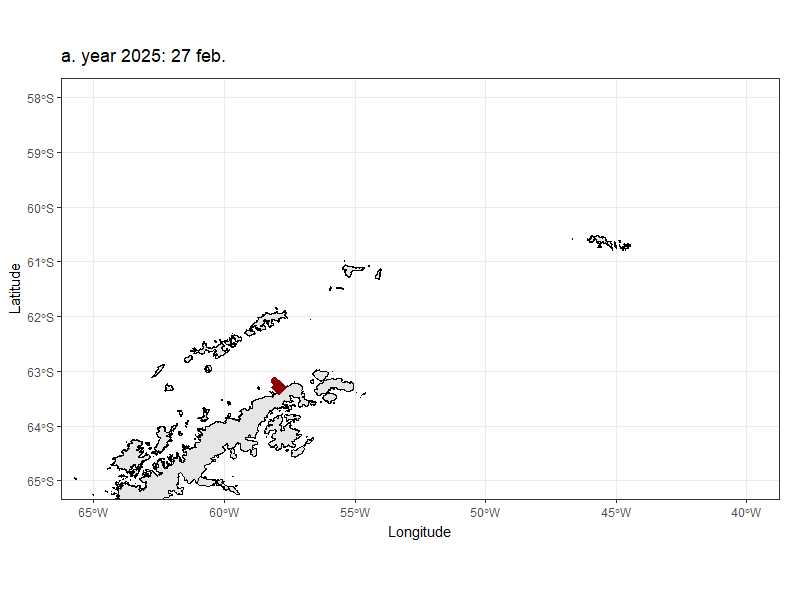

### supplementary video S3

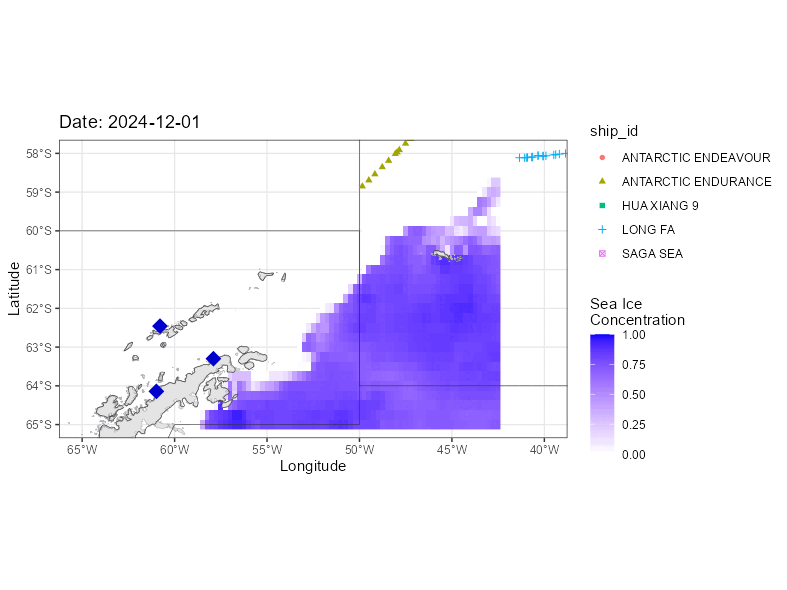
